# Multiscale Bayesian Simulations Reveal Functional Chromatin Condensation of Gene Loci

**DOI:** 10.1101/2023.12.01.569511

**Authors:** Giovanni B Brandani, Chenyang Gu, Soundhararajan Gopi, Shoji Takada

**Affiliations:** Department of Biophysics, Graduate School of Science, Kyoto University, Kyoto 606-8502, Japan

## Abstract

Chromatin, the complex assembly of DNA and associated proteins, plays a pivotal role in orchestrating various genomic functions. To aid our understanding of the principles underlying chromatin organization, we introduce Hi-C metainference, a Bayesian approach that integrates Hi-C contact frequencies into multiscale prior models of chromatin. This approach combines both bottom-up (the physics-based prior) and top-down (the data-driven posterior) strategies to characterize the 3D organization of a target genomic locus. We first demonstrate the capability of this method to accurately reconstruct the structural ensemble and the dynamics of a system from contact information. We then apply the approach to investigate the Sox2, Pou5f1, and Nanog loci of mouse embryonic stem cells using a bottom-up chromatin model at 1kb resolution. We observe that the studied loci are conformationally heterogeneous and organized as crumpled globules, favoring contacts between distant enhancers and promoters. Using nucleosome-resolution simulations, we then reveal how the Nanog gene is functionally organized across the multiple scales of chromatin. At the local level, diverse chromatin folding motifs correlate with epigenetics, with open chromatin predominantly observed at cis-regulatory elements and compact tetranucleosomes in between. At the larger scale, we find that enhancer-promoter contacts are driven by the transient condensation of chromatin into compact domains stabilized by extensive inter-nucleosome interactions. Overall, this work highlights the condensed, but dynamic nature of chromatin *in vivo*, contributing to a deeper understanding of gene structure-function relationships.

## Introduction

Chromatin, the complex of DNA and proteins organizing the Eukaryotic genome, plays a pivotal role in orchestrating a multitude of genomic functions, including DNA replication (1) and the precise regulation of gene activity (2). Its structure exhibits a hierarchical organization, with biological function affected by structural changes across all levels of the hierarchy (3). The fundamental unit of chromatin is the nucleosome, an assembly comprising of about 150 base pairs (bp) of DNA wrapped around a histone octamer (4). Nucleosomes are not simply barriers preventing access to genomic DNA; rather, they can be better understood as flexible platforms on top of which key biological processes can be organized (5). For example, nucleosomal DNA sliding and unwrapping from histones modulate transcription factor (TF) binding (6–8), with downstream effects on gene expression. At a higher hierarchical level, these nucleosomes interact together to form chromatin fibers exhibiting remarkable structural polymorphism (9–11). *In vivo*, these fibers are predominantly disordered (12), but specific chromatin motifs are associated with distinct genomic regions, such as gene promoters or gene bodies (13). Chromatin fibers further fold into topologically associated domains (TADs), contiguous regions with enriched physical contacts separated by insulator elements (14). The pattern of TADs along the genome influences the communication between cis-regulatory elements (CREs) such as enhancers and promoters (15, 16). Pointing to the intimate link between genome structure and function, genetic engineering has shown a correlation between the probability of enhancer-promoter contacts and gene expression levels (17), while transcription bursting dynamics was found to be linked with enhancer-promoter looping over time (18).

Despite substantial progress in recent years, our understanding of the physical principles governing chromatin organization across different scales of genes remains limited, which hinders further exploration of the interplay between genomic structure and function. In this work, we integrate *in vivo* experimental data into polymer models to explore the 3D structure of mammalian genes from the local chromatin fiber level to long-distance interactions between CREs. Our findings shed light on how this organization can contribute to gene regulatory functions.

Theoretical approaches rooted in polymer physics have become increasingly important for complementing experiments and uncovering genomic structure-function relationships (19). Some of these approaches are data-driven (20–23), with interactions between genomic sites optimized to match the experimental Hi-C contact frequencies (24). Others are centered around a key mechanism responsible for 3D genome organization, such as SMC-driven loop extrusion (25), protein-DNA bridging (26), or phase separation due to epigenetic modifications (27, 28). Furthermore, advances in the development of bottom-up molecular models of chromatin (29, 30), parametrized based on the physics of its biomolecular components, now offer a viable route to provide a mechanistic understanding of genes based on first principles (31). Such models have revealed some of the molecular determinants of chromatin organization, including the role of nucleosome plasticity in mediating liquid-liquid phase separation (32). Notably, Tamar Schlick and co-workers utilized a nucleosome-resolution model to explore the structure of the HOXC gene locus in mouse embryonic stem cells (mESCs) (33), identifying the spontaneous formation of loops linking gene promoters with other regions of the locus. However, such physics-based chromatin models typically do not capture the effects of all the factors constituting the complex cellular environment, such as potential effects originating from TF binding (26, 34). Due to the practical consideration in modeling the behavior of several contributing molecular factors, physics-based models may not always accurately reproduce the experimental organization of chromatin *in vivo*, as determined from Hi-C or Micro-C contact probabilities (24, 35).

To address this limitation, Bayesian methods offer a framework to incorporate experimental observations into an existing physical model (36, 37). Metainference is one such Bayesian approach originally developed to integrate NMR data into all-atom force fields for protein molecular dynamics (MD) simulations (37). We introduce and validate Hi-C metainference, an extension of the original approach to augment an existing prior model of chromatin to account for the observed *in vivo* contact probabilities of a target locus (from Hi-C (24), Micro-C (35), or other related approaches (13)), allowing one to characterize its 3D organization by MD simulations. As priors, we employ two distinct bottom-up coarse-grained models of chromatin: the 1CPN model (30) at nucleosome resolution, previously parametrized from near-atomistic nucleosome simulations (38), and a chromatin polymer model at 1kb resolution newly parametrized from extensive 1CPN simulations. Therefore, our exploration of the 3D genome organization can be regarded as a 2-way process where the bottom-up prior is adjusted in a top-down fashion by integrating the experimental contact frequencies, retaining the advantages of both physics-based and data-driven approaches.

In this study, we integrate Micro-C data (39) into our chromatin models and investigate the organization of the Sox2, Pou5f1, and Nanog genomic loci in mESCs. First, our simulations show that *in vivo* genes adopt relatively compact conformations, with the same size scaling of a crumpled globule (40), while at the same time displaying significant structural heterogeneity and fast dynamics. Using a nucleosome-resolution prior chromatin model for a section of the Nanog locus, we find that the local organization of the chromatin fiber is highly dependent on the underlying epigenetic marks, being open at CREs and more compact elsewhere. Finally, we demonstrate that transient, direct contacts between the Nanog promoter and its nearest enhancer are driven by the condensation of the chromatin fiber into a compact domain through extensive nucleosome-nucleosome interactions. Our simulations suggest that chromatin plasticity over multiple length-scales provides a physical basis for essential gene regulatory functions.

## Results

### Bayesian integration of Hi-C data into prior physics-based chromatin models

We introduce Hi-C metainference to update an existing prior model of chromatin (Fig. 1A, e.g., nucleosome-resolution 1CPN (30) or coarser 1kb-resolution model) using *in vivo* genomic contact frequencies from experiments such as Hi-C (24) and Micro-C (35). Within this framework, MD simulations are used to sample the conformational ensemble of the target system from the posterior probability distribution according to the Bayes’ theorem (37). The infinite ensemble is represented using a finite number N of replicas of the same chromatin system (Fig. 1A, top left panel, for N=4) – for example a polymer made of L beads, each representing a certain number of base pairs of genomic DNA. The use of replicas makes metainference ideally suited to model ensemble-averaged experimental data produced by highly heterogenous systems such as chromatin (41). Given a chromatin structure, we use a forward model that computes the LxL contact map of each replica (Fig. 1A, bottom left panel). We then average the contact maps over the N replicas to compute a map of contact probabilities, which can be compared to the contact frequencies measured in experiments (Fig. 1A, bottom right panel). The dynamics of the N replicas is then simulated in parallel using the interaction potential of the prior chromatin model together with an additional Bayesian energy score accounting for the mismatch between experiments and predictions from the MD ensemble. This extra bias represents *in vivo* effects originating from epigenetics (21, 27, 42) and TF binding (26, 34) that are not explicitly included into the physics-based model. These steps are repeated each MD timestep until we obtain a converged conformational ensemble (see Methods for more details).

**Figure 1:**
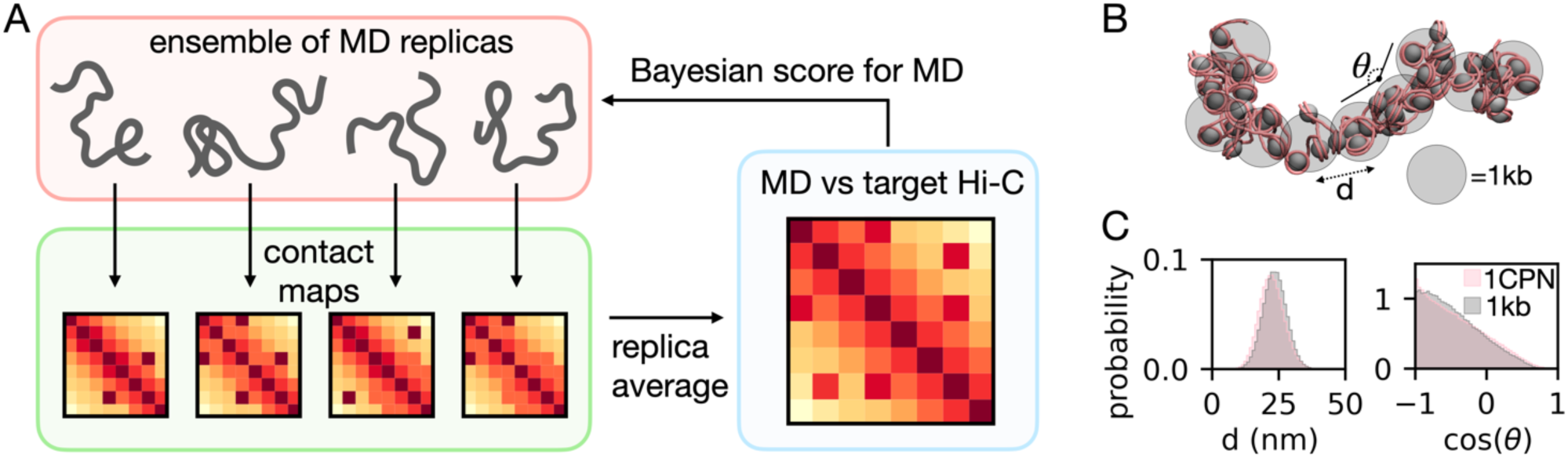
Hi-C metainference approach and prior models. A. In Hi-C metainference, we simulate by MD a target genomic region using N replicas (N=4 in example), representing the (in principle infinite) conformational ensemble of the system (top panel). For each conformation, we generate a contact map using a forward model (bottom left). We average the contact maps over the N replicas to generate a map of contact probabilities to be compared with the target Hi-C map (bottom right). To integrate the Hi-C data into the simulation, we add to the interaction potential of the prior model a Bayesian energy score based on the difference between the MD and the target Hi-C map. B. Representations of the nucleosome-resolution 1CPN and 1kb prior chromatin models, indicating the 1kb model bonds and angles whose potentials are parametrized based on the 1CPN simulations. In the 1CPN structure, nucleosomal DNA is represented in pink, while the histone octamers are represented as dark gray particles. In the 1kb model, each coarse-grained bead (in light gray) represents 5 nucleosomes. C. Comparison of the probability distributions of the bond distances (left) and angles (right) between 1kb coarse-grained particles from the 1kb-resolution (gray) and the 1CPN (pink) chromatin models.

In principle, Hi-C metainference can be applied using any prior MD model of chromatin, as different models cater to the needs of different applications. In this work, we use two chromatin prior models at different resolution: the nucleosome-resolution 1CPN chromatin model (30), and a new coarse-grained 1kb-resolution model parametrized from 1CPN. The 1CPN model has been rigorously parametrized using a bottom-up procedure based on simulations of nucleosome-nucleosome interactions at near-atomic resolution (38), and validated against experimental chromatin sedimentation coefficients. In the 1kb model, chromatin is represented as a polymer consisting of 1kb beads (=5 nucleosomes) connected by bond and angle potentials (Fig. 1B, see Methods for more details). The interaction parameters are optimized in a bottom-up way to match the distributions of bond distances and angles from long 1CPN simulations of 50-nucleosome fibers with a nucleosome repeat length of 200bp (Fig. 1C and Fig. S1).

### Validation by Copolymer Model

In order to test the capability of metainference to reconstruct a target polymer ensemble from its Hi-C map, we setup a test using a 200 kb copolymer made of (L=200) 1kb beads modeled as the prior chromatin model, except that three 10kb regions (E1, E2, E3) act as sticky CREs with relatively strong interactions with each other and weaker interactions with the rest of the locus (gene bodies-like). This model produces a complex copolymer ensemble with a variety of interactions between the CREs, where these can be paired in the three possible combinations, or all be far away from each other, or all collapse into one single hub (Fig. 2A, top panel). We test whether metainference using the non-informative prior chromatin model and the copolymer contact map can reconstruct both the statistics and the dynamics of the target copolymer.

**Figure 2:**
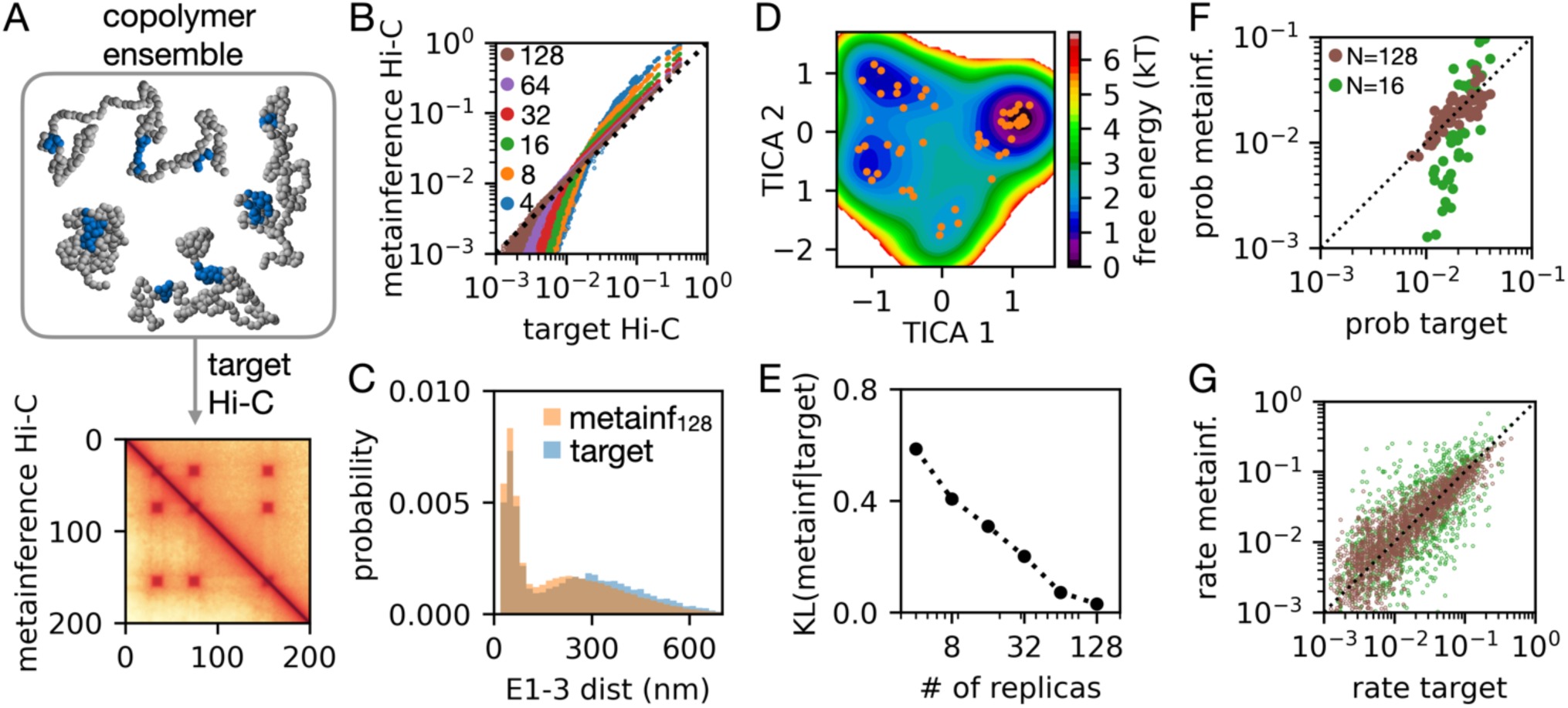
Hi-C metainference validation on complex copolymer ensemble. A. Target Hi-C map (right upper half in the bottom plot) generated by simulations of a 200kb copolymer containing three sticky CREs (in blue in representative snapshots on top), vs the Hi-C map generated from metainference (left lower half) simulations using N=128 replicas and the prior 1kb chromatin model. B. The correlation between the Hi-C contact probabilities estimated from metainference MD (y-axis) and from the original copolymer ensemble (x-axis). Different colors correspond to different numbers of replicas. C. Comparison of the probability distributions of the distances between the CREs E1 and E3 obtained from the target copolymer ensemble (blue) and metainference MD with N=128 replicas (orange). D. Free energy landscape of the N=128 metainference simulations projected on the first two TICA coordinates, showing the 50 cluster centers in orange. E. Kullback-Leibler (KL) divergence of the metainference cluster probability distribution from the target copolymer distribution, as a function of the number of replicas. F-G. Comparison of the cluster equilibrium probabilities (F) and MSM-estimated transition rates (G) between the metainference MD (y-axis) and the target copolymer system (x-axis) for either N=16 (green) or N=128 (brown) replicas.

Figure 2A shows the good agreement between the target contact map and the one generated from the metainference simulations using N=128 replicas. The correlation between the contact probabilities increases as the number of replicas in increased from 4 to 128 (Fig. 2B). This is to be expected, since reproducing the formation of a contact with probability p requires roughly N=1/p or more replicas, as the contact should appear at least once in the ensemble of replicas used to represent the true infinite ensemble during the MD simulation. Hi-C metainference can also recover properties not directly used as a bias, such as the distribution of distances between the different CREs (Fig. 2C, Fig. S2A,B) and radius of gyration (Fig. S2C). To provide a rigorous comparison of the ensemble statistics, we divided the conformations observed in the copolymer and metainference simulations into 50 clusters using k-means (43) after performing a TICA dimensionality reduction (44) (see Supporting Information (SI) Methods). First, the free energy landscape of the N=128 metainference system projected on the first two TICA coordinates (Fig. 2D) is close to the one obtained from the target copolymer simulations (Fig. S2D). The equilibrium probabilities of the different clusters are in good agreement (Fig. 2F), and the agreement, as measured by the Kullback-Leibler divergence between the distributions, increases with the number of replicas used (Fig. 2E). Finally, we also confirmed that metainference can also recover the dynamics of the target copolymer system, as shown by the good correlation between the cluster transition probabilities estimated by Markov state modeling (45) of the metainference and target copolymer simulations (Fig. 2G, see SI Methods for details of the Markov state modeling).

### mESCs chromatin has a compact but heterogenous 3D organization

We next applied Hi-C metainference to explore the 3D organization of the 300kb Pou5f1 and Sox2 genomic loci of mESCs by integrating the corresponding Micro-C data (39) into our 1kb chromatin model. Metainference using 128 replicas produces Hi-C maps in good agreement with the experiments (Fig. 3A,B). In Fig. 3C and 3D, we compared the distributions of the physical distances between the promoters and nearby genomic locations with single cell measurements made when the genes are actively expressed (46), finding that both averages and standard deviations are in good agreement with these experiments. These results suggest that the transcription status of the genes have a minor impact on the experimentally measured enhancer-promoter distances at these loci, consistent with a recent analysis (47).

**Figure 3:**
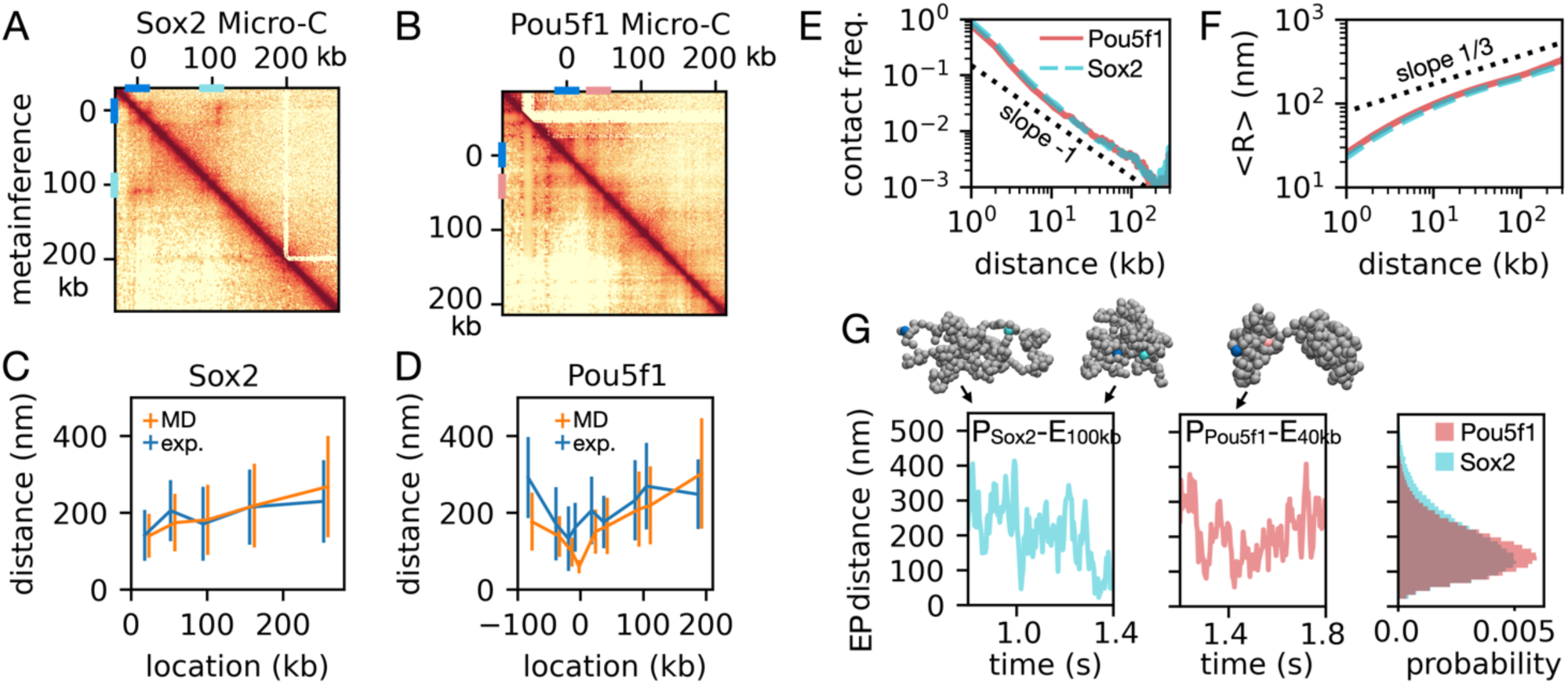
Structural heterogeneity of the Pou5f1 and Soc2 loci of mESCs. A and B. Comparison of Micro-C contact maps of the 300kb loci of the Pou5f1 (A) and Sox2 (B) genes with the corresponding results obtained from metainference with N=128 replicas. The genomic locations of the two promoters (0kb), the Sox2 +100kb enhancer and the Pou5f1 +40kb enhancer are indicated on the contact maps as blue, cyan, and pink bars, respectively. C and D. Comparison of the averages and standard deviations (size of the bars) of the distributions of physical distances between the Pou5f1 (C) and Sox2 (D) promoters (0kb) with nearby genomic locations obtained from single molecule experiments (46) (blue line) and from metainference MD based on the Micro-C maps (orange). E-F. Scaling of contact frequency (E) and average end-to-end distance <R> from metainference MD as a function of genomic distance (Pou5f1, red solid line; Sox2, cyan dashed line). Except for very short distances L, scaling is close to that of a crumpled globule (40), with contact frequency∼1/L and <R>∼L^1/3^. G. Representative trajectories and snapshots of the Sox2 (left) and Pou5f1 (center) genomic loci, and the corresponding enhancer-promoter (EP) distance distributions (right) computed from the metainference MD ensembles (colors as in E-F). In the simulation snapshots, promoters are highlighted in blue, while the Sox2 enhancer at +100kb and the Pou5f1 enhancer at +40kb are in cyan and pink, respectively.

Except for very small genomic distances L, we find that the studied genomic loci are spatially organized as a crumpled globule (40), with the frequency of contacts scaling as ∼1/L (Fig. 3E, as over very large distances (24)) and the average end-to-end distances scaling as <R>∼L^1/3^ (Fig. 3F). Such scaling was also found in a recent *in vivo* microscopy study of a Drosophila gene (18), highlighting how genes adopt relatively compact conformations favoring the formation of direct contacts between distant CREs. Despite such compact scaling, chromatin explores very heterogeneous conformational ensembles, sampling not only globular domains characterized by extensive contacts between distant regions, but also more open conformations with fewer contacts (see snapshots in Fig. 3G). As indicated by the rapid decay of the enhancer-promoter distance autocorrelation functions (Fig. S3A), these loci typically undergo fast transitions between conformations with small (<50nm) and large (>300nm) enhancer-promoter distances (Fig. 3G), over a timescale of less than a second, typically much faster than the timescales of gene bursting dynamics (48). Interestingly, the distributions of enhancer-promoter distances peak around ∼150 nm (Fig. 3G, left), as the size of the condensed but liquid domains experimentally observed in active chromatin (49). This suggests an active role of such liquid domains in CRE communication.

We then applied Hi-C metainference to study the structural ensemble and the kinetics of the 200kb Nanog locus of mESCs. Nanog gene expression is regulated by three enhancers, located at –45kb (E1), –5kb (E2), and +60kb (E3) relative to the promoter (50). This locus represents a model system to explore the coupling between 3D gene organization and function, and in particular the potential co-operativity of the three enhancers. Fig. 4A shows the good agreement between locus contact frequencies from Micro-C (39) and from the metainference simulations obtained using N=128 replicas. As in the cases of the Sox2 and Pou5f1 loci, the Nanog locus is characterized by a heterogeneous conformational ensemble, with dynamic domains bounded at variable locations (Fig. 4B). Reflecting this heterogeneity, the enhancers sample a wide variety of distances to the promoter (Fig. S4A), sometimes even <50nm, indicating the possibility of direct physical contacts. Notably, all CREs often cluster together into a single hub (Fig. 4B, bottom left snapshot, and Fig. S4B for the free energy landscape projected on a pair of enhancer-promoter distances), suggesting that the three enhancers may co-operatively activate their target (51), or compensate for each other’s loss (50). As the other loci, the dynamics of the Nanog gene is fast, with the system able to explore highly diverse configurations on the second timescale (distance autocorrelations in Fig. S4C).

**Figure 4:**
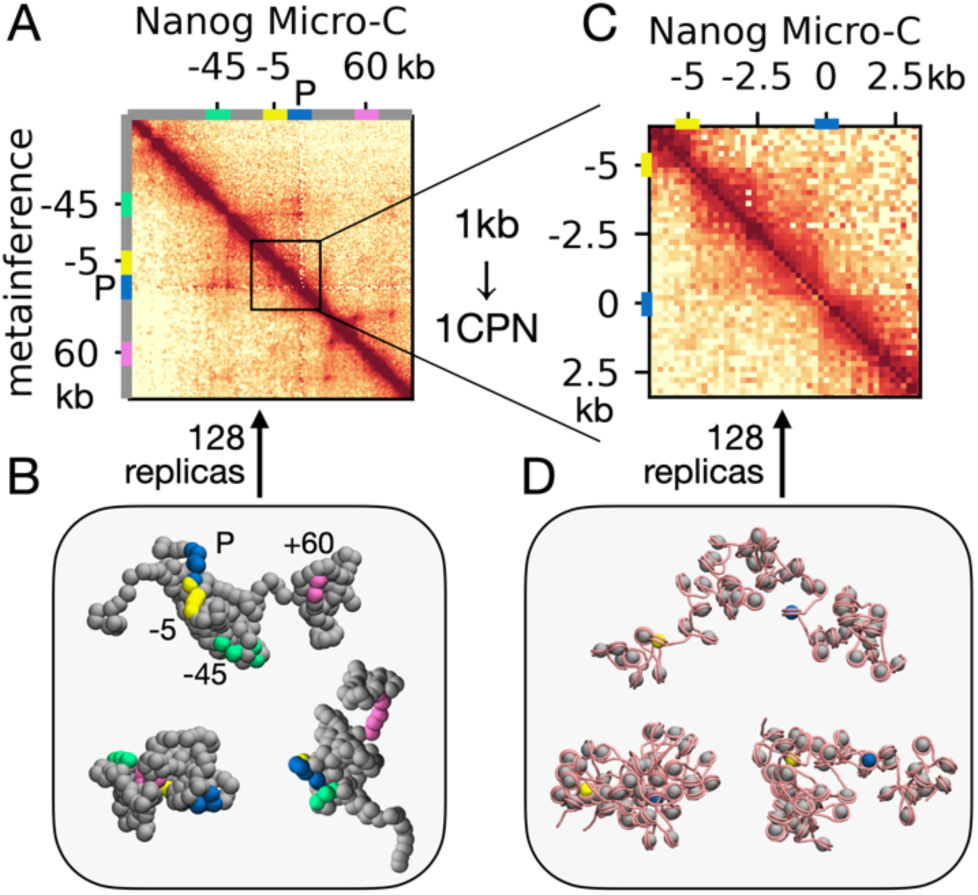
Multiscale Nanog locus organization. A. Comparison of the 200kb Nanog locus Micro-C map as observed in mESCs with the result form metainference using N=128 replicas and the 1kb model. The locations of the Nanog promoter and the –45kb, –5kb and +60kb enhancers are indicated on the contact map axes with the blue, green, yellow, and pink bars, respectively. B. Snapshots of representative Nanog conformations. CRE locations are highlighted with the same color scheme as in A. C Comparison of Micro-C and 128-replica metainference contact maps applying the 1CPN chromatin model to the 10kb locus including the Nanog promoter (blue bar) and its closest enhancer 5kb upstream (yellow). D. Representative 1CPN conformations of the 10kb locus sampled by metainference.

To gain insights into the finer organization of the Nanog locus at nucleosome resolution, we ran further 1CPN (30) metainference simulations of the 10kb region encompassing the promoter and its closest enhancer located 5kb upstream. As in the 1kb model, we find a good agreement between the metainference and reference Micro-C map at 200bp resolution (Fig. 4C), while the ensemble of chromatin conformations (further discussed in the next section), is also significantly heterogenous (Fig. 4D).

### Nanog organization at nucleosome resolution reveals functional chromatin condensation at local and global scales

From the 1CPN metainference simulations, we first explored the local organization of the Nanog locus chromatin fibers. We focused our analysis on tetranucleosomes as the fundamental units of the fiber organization (9–11, 13). We performed principal component analysis (PCA) (43) of all tetranucleosome contact maps within the 10kb Nanog locus in all the 128 metainference replicas. The free energy landscape (logarithm of probability distribution) along the tetranucleosome projections on PCA components 1 and 3 highlights distinct populations of structures stabilized by different inter-nucleosome contacts (Fig. 5A, left panel, 7 clusters identified by K-means clustering (43) based on their contact maps, 4 of them considered for further analysis). PCA 1 represents roughly the overall number of inter-nucleosome interactions, while PCA 3 represents the difference between i,i+2 contacts and other (i,i+1 and i,i+3) contacts (PCA 2 is not symmetric under a change in DNA direction, so it has been excluded from the analysis, Fig. S5). Notably, the landscape includes significant populations of tetranucleosomes resembling experimentally determined structures stabilized by either i,i+1 (9) (Fig. 5A, bottom right) or i,i+2 (10, 11) (Fig. 5A, top right) stacking interactions, which if propagated would lead to the formation of regular 1-start and 2-start chromatin fibers (55), respectively. However, the most populated structures in the landscape are actually stabilized by a mixture of i,i+1 and i,i+2 interactions, indicative of an overall disordered chromatin fiber organization (as can be appreciated also from the representative Nanog locus conformations in Fig. 4D). Interestingly, we also find a population of compact tetranucleosomes with all-to-all direct interactions, and another population of very open tetranucleosomes completely lacking inter-nucleosome contacts (Fig. 5A, two middle panels on the right). As suggested based on Hi-CO experiments on yeast (13), local fiber organization correlates with chromatin epigenetic features. We find that the open tetranucleosomes are prevalently observed at the CREs of the locus, as determined from the ChIP-seq profiles of RNAPII binding (52) and H3K27ac tail modification (53) (Fig. 5B). Such open chromatin at CREs may facilitate TF and RNAPII binding, which in turn could lead to the formation of nucleosome free regions or fragile nucleosomes further disrupting the fiber (56). On the contrary, compact tetranucleosomes are prevalently found outside CREs (Fig. 5B). Such compact structures may facilitate the communication between CREs. Analysis of the average distances between any given nucleosome and the nearest one in space confirms that chromatin is more open at CREs and more compact in between (Fig. S5B). The average nearest nucleosome distance throughout the locus is ∼12 nm, consistent with the value obtained in our prior 1CPN simulations and the experimental value obtained by *in situ* cryo-ET of human T-lymphoblasts interphase chromatin (12).

**Figure 5.**
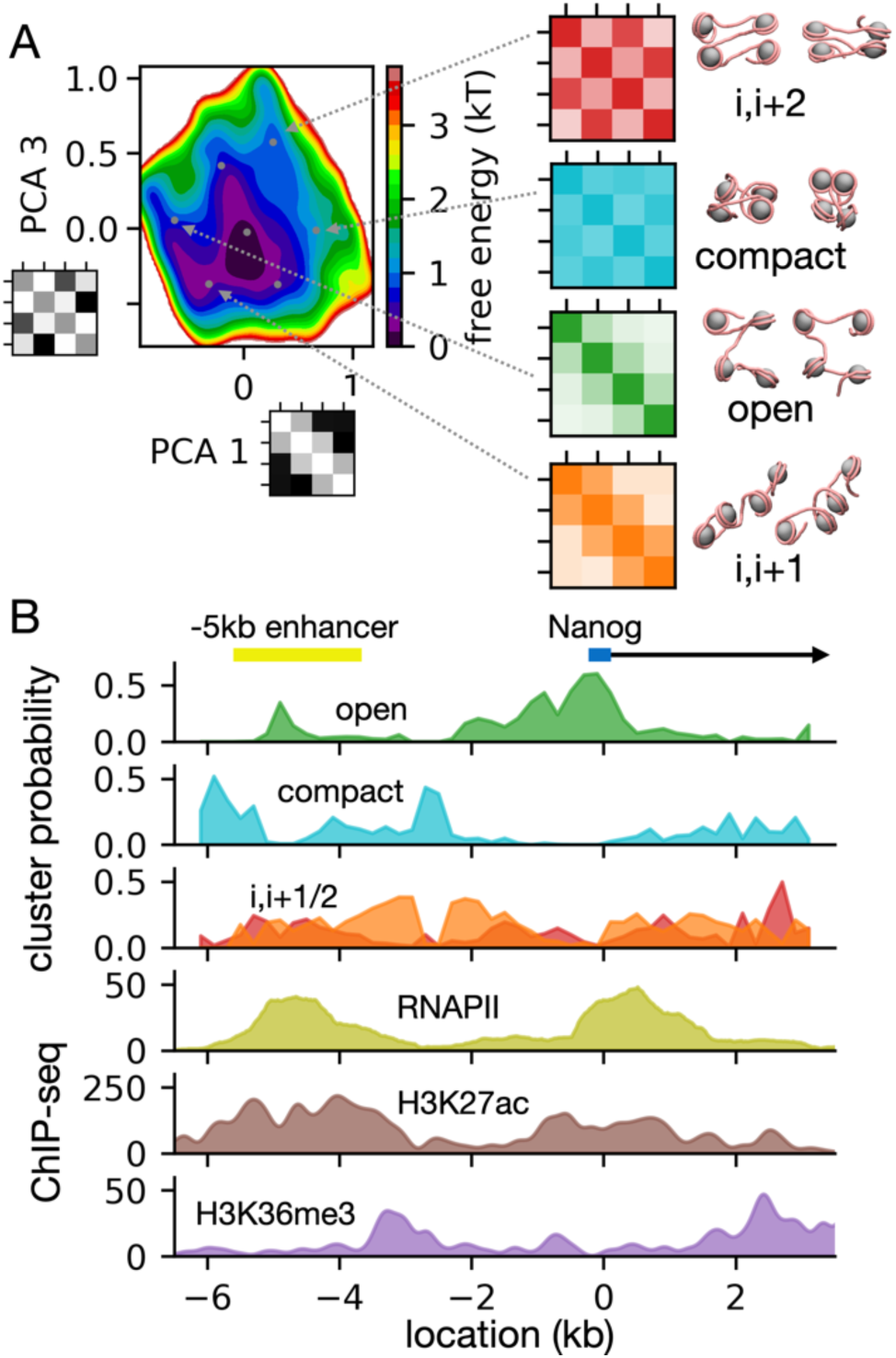
Nanog local chromatin organization. A. Principal component analysis (PCA) (43) of the 10kb Nanog locus tetranucleosome structures based on their contact maps. The left panel shows the free energy landscape along the 1^st^ and 3^rd^ PCA components (the maps next to the labels indicate the component eigenvectors). PCA 1 captures the total number of contacts, while PCA 3 correlates with the difference between i,i+2 and other (i,i+1 or i,i+3) contacts. The landscape shows 7 k-means clusters (43) (cluster centers in gray) stabilized by different nucleosome-nucleosome interactions. On the left we highlight contact maps and representative structures of 4 tetranucleosome clusters, characterized by regular i,i+2 contacts (red map), compact structure (cyan), open structure (green), or regular i,i+1 contacts (orange). B. Top three panels: probabilities of the 4 tetranucleosome clusters highlighted in panel A as a function of genomic location within the 10kb Nanog locus. Three bottom panels: the experimental ChIP-seq signals of RNAPII binding (GEO accession: GSE22562 (52)), and H3K27ac (GEO accession: GSM2417096 (53)) and H3K36me3 (GEO accession: GSM2417108 (53)) epigenetic modifications. Chromatin is relatively open at CREs, while compact elsewhere. On top of the graphs, we also indicate the locations of the Nanog promoter (blue bar) and the –5kb enhancer region (yellow bar) (54).

How does chromatin structure support the formation of direct contacts between promoter and –5kb enhancer? To discuss this question, we performed PCA dimensionality reduction of the complete 10kb chromatin contact maps observed in the 1CPN metainference simulations (Fig. 6A). K-means clustering using the top 4 PCA components and visual inspection of the PCA projection highlights 4 clusters characterized by distinct enhancer-promoter distances (Fig. 6B) and chromatin organization (Fig. 6C). The respective cluster-averaged contact maps and representative conformations (Fig. 6C) provide insights into the organization: in the extended cluster (58% probability), nucleosomes mainly interact with their closest neighbors and chromatin is relatively open; in the promoter (P) domain cluster (15%) there are some contacts between the promoter and the gene body downstream from it; the enhancer-promoter (EP) domain cluster (21%) is characterized by a distinct compact domain bounded by the –5kb enhancer upstream and the Nanog promoter downstream; the loop cluster (6%) has a hairpin-like organization where chromatin adopts an overall condensed configuration with extensive inter-nucleosome interactions between distant regions. In the extended and P domain clusters, the physical distances between promoter and enhancers are typically between 50 and 100 nm, while in the EP domain and loop clusters they are less than 50 nm (Fig. 6B), sufficient for direct contacts mediated by the RNAPII-mediator complex (57). We note how the global organization of EP domain mirrors the folding of tetranucleosomes at the lower scale: with open chromatin motifs at the CREs establishing the boundaries of a central domain rich in compact tetranucleosomes. The condensed nature of EP domain and loop conformations is well illustrated by the fact that the average distances from the promoter do not significantly depend on genomic distance (Fig. 6D). These data reveal the high plasticity of chromatin within the Nanog locus, with predominant extended conformations that transiently condense into compact domains supporting direct contacts between enhancer and promoter (Fig. 6E). Notably, such contact formation is also observed in our unbiased 1CPN simulations, where reversible chromatin condensation events occur on a sub-ms timescale (Fig. S6). This suggests that the physical principles underlying the folding of chromatin fibers, essentially nucleosome plasticity and nucleosome-nucleosome interactions, may be sufficient to spatially and dynamically organize the 10kb Nanog locus, without requiring additional forces from other DNA binding proteins.

**Figure 6.**
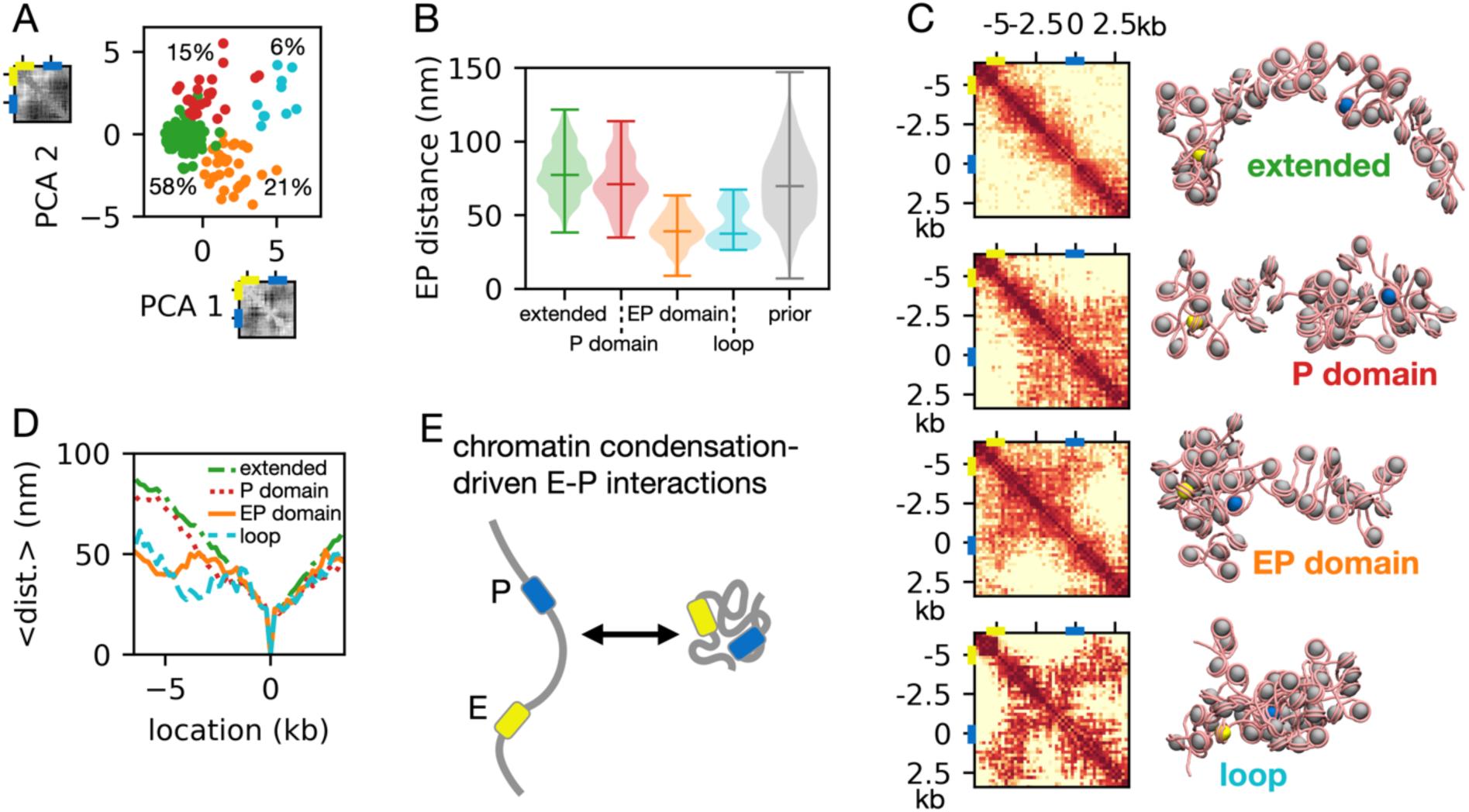
Global organization of the 10kb Nanog locus at nucleosome resolution. A. Projection of the 10kb Nanog locus chromatin conformations along the first two PCA coordinates obtained from their global contact maps. The component eigenvectors are shown next to the respective labels, indicating that PCA 1 roughly correlates with the total number of contacts, while PCA 2 with the difference between contacts in the right and left side of the locus. The projection reveals 4 k-means clusters characterized by different global 3D organization (the clusters are indicated by different colors, together with their population). B. Distributions of enhancer-promoter (EP) distances across the 4 clusters highlighted in panel A, together with the overall distribution for the unbiased 1CPN simulations. The bars in the violin plots indicate the medians and the extrema of the distributions. The 4 clusters are referred to as extended (green), promoter (P) domain (red), enhancer-promoter (EP) domain (orange), and loop (cyan). The distribution obtained from the prior 1CPN simulations is shown in gray. C. Population-averaged contact maps and representative structures of the 4 clusters. The promoter and enhancer locations are indicated by the blue and the yellow nucleosomes, respectively. D. Average contact frequencies and average distances to the promoter for the 4 clusters, highlighting how the EP domain and loop clusters represent condensed conformations where the promoter forms direct contacts with other locations regardless of the genomic distance. F. Schematics highlighting that direct enhancer-promoter contacts are driven by the condensation of the chromatin structure into a compact globule.

## Discussion

In this study, we demonstrated the successful application of metainference to integrate Hi-C contact frequencies into MD simulations of chromatin. Within this Bayesian scheme, the original polymer model of chromatin constitutes our prior, while the polymer conformations are sampled from the posterior probability distribution accounting for the contact frequencies observed in the experimental Hi-C map. This makes our approach both bottom-up, starting from a physics-based prior suitable for exploring molecular mechanisms, and top-down, providing a picture of 3D genomic organization consistent with experimental data. Our tests showed that Hi-C metainference can reconstruct both the structural ensemble and the dynamics of a target system based on contact information, with accuracy increasing with the number of replicas used to model the (in principle infinite) ensemble. A related Bayesian approach to the 3D genome (36) requires comparing the results of multiple simulations using different numbers of replicas to find the optimal minimal ensemble representing the target system. Instead, metainference is valid for any number of replicas N>1, although in practice we recommend setting this to the inverse of the minimum contact probability that one aims at capturing in the final 3D ensemble (N=128 gives a reasonable value for our applications). We note that while here we solely focused on applications to genome organization, the generality of this approach makes it applicable to other situations where we may want to integrate contact information into a prior molecular model. For instance, protein cross-linking data may be used to reconstruct the structure of large molecular complexes (58).

Metainference is particularly convenient for its flexibility, in that one can integrate Hi-C data into any chromatin model suitable for the target application. As previously shown (37), an accurate prior molecular model can help getting closer to the true conformational ensemble of the system, and therefore producing realistic representations of the genome likely benefits from the use of physics-based models rigorously parametrized based on the interactions between the fundamental components of chromatin. By adding an extra bias on top of an existing prior model, metainference can effectively account for *in vivo* factors that are not directly represented, such as the binding of TFs (26, 34) or epigenetic modifications (21, 27, 33, 42). Alternatively, the differences in the conformational ensembles obtained using the physics-based models with and without metainference can guide the modeling of necessary additional factors into the prior model or suggest hypotheses on the underlying mechanism. For the applications to the genomic organization of mESCs presented here, we integrated Micro-C data (39) into two physics-based models of chromatin, the nucleosome-resolution 1CPN model (30) and a newly parametrized 1kb model, although other models may also be employed (29, 59).

The interplay between the 3D genome organization and the regulation of gene expression is a fundamental problem in molecular biology, for which a clear mechanistic understanding remains elusive. As observed from super-resolution chromatin tracing experiments (41) and other polymer modeling approaches (26, 36), we confirm that functional genomic interactions occur within the context of a highly heterogeneous conformational landscape that includes both open and compact conformations. In the studied loci, we find frequent enhancer-promoter contacts separated by a short distance on the order of the mediator-RNA polymerase complex (∼50nm) (57), indicating their potential to directly control gene expression based on proposed models of enhancer action (60). At the Nanog locus, the chromatin often collapses into a compact structure where all three enhancers are at close distance to the Nanog promoter, possibly representing a co-operative transcription hub (51). Overall, the studied loci are organized as crumpled globules (40), with hierarchically formed globular domains encompassing increasing genomic lengths. Such fractal-like organization was already suggested based on Hi-C data (24), and further confirmed by a recent microscopy study in Drosophila (18), where it was shown to promote the communication of distant CREs.

Past research offers contradictory evidence (61, 62) on the importance of 3D genome structure for gene regulation, with various mechanistic models of enhancer activity still being currently debated (63). Issues may be partially due to the nonlinearity of enhancer-promoter communication (17, 64), with contact frequencies affecting gene expression levels only within a narrow range of values. This nonlinearity has been suggested to originate from a separation of timescales between genomic structural fluctuations and gene activation (17). Indeed, consistently with recent simulations (59), we find that the Sox2, Pou5f1, and Nanog loci of mESCs can rapidly explore their vast space of 3D conformations. Transient contacts between CREs (potentially stimulating gene expression) are formed and broken on a timescale of less than a second, significantly faster than the timescales of gene activation, on the order of several minutes (48). This separation of timescales helps explain difficulties in the experimental detection of correlations between 3D genome structure and gene expression (47).

Our 1CPN simulations of the Nanog locus offer an opportunity to explore how the detailed molecular structure of chromatin can support the functional organization of genes across multiple length-scales. We find that the extent of inter-nucleosome interactions critically shapes both the local and the global folding of the chromatin fiber, leading to a surprising conformational plasticity. At the local level, chromatin appears relatively disordered, consistent with experimental observations *in situ* (12), but with the occasional formation of regular tetranucleosome motifs resembling those characterized in past in vitro studies (9–11). Importantly, we find that open chromatin motifs lacking inter-nucleosome contacts are predominantly observed at CREs. Here, specific epigenetic modifications such as histone tail acetylation promote the opening of chromatin (65, 66), which in turn should facilitate the recruitment of TFs and RNA polymerases and contribute to further opening (56). On the other hand, compact tetranucleosome motifs are enriched in between CREs, where they may help drive these elements in closer spatial proximity. Such kind of epigenetic-dependent local chromatin organization was previously suggested in yeast based on simulations and Hi-CO experimental contact frequencies (13). However, the tetranucleosome landscape observed at the Nanog locus appears richer than the binary set of alpha and beta motifs in yeast, perhaps thanks to the longer DNA linker length in mammalian chromatin (67). We note that our metainference simulations could capture the role of epigenetics despite the use of a uniform prior model of chromatin, confirming the successful integration of experimental contact frequencies to account for key factors missing in the model.

At the global level, classic models of Eukaryotic gene regulation depict enhancer-promoter communication as based on the formation of a loop linking together the two CREs (63). However, our 10kb Nanog locus simulations suggest that communication relies on dynamic transitions between extended chromatin fibers and condensed domains characterized by direct contacts between CREs. Interestingly, such plasticity is also observed in 1CPN simulations of chromatin lacking any experimental bias. Over large genomic distances, the 3D genome is often organized by proteins through processes such as SMC-driven loop extrusion (25, 68) and bridging interactions (26, 69). Instead, we suggest that inter-nucleosome interactions alone are sufficient to explain the observed dynamic condensation of the 10kb Nanog locus. Despite these general principles, the detailed organization of each locus will be shaped by the local epigenetics. Overall, the 10kb Nanog locus appears to be hierarchically organized so that chromatin folding at the local level orchestrates folding over larger scales. This is particularly observed in a large population of structures with direct enhancer-promoter contacts stabilized by the formation of a globular domain rich in compact tetranucleosomes bounded by open chromatin at the CREs (EP domain cluster in Fig. 6).

Overall, we showed how Hi-C metainference allows the efficient integration of experimental data and physical modeling to explore the mechanisms underlying the multiscale spatial organization of the genome. Our simulations reveal a condensed, liquid-like chromatin whose structure and dynamics is ideally suited to support genomic function, in line with the picture recently emerging from experiments (49, 70, 71). Ultimately, our integration of theoretical modeling and experimental data will aid our understanding of the physical basis of gene regulation (60) and other key genomic functions.

## Materials and Methods

### Hi-C Metainference Theory

Given the experimental contact frequency f that two genomic loci are physically close in an ensemble of cells, the posterior probability distribution of an ensemble of chromatin conformations 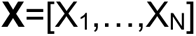 is proportional to (37):

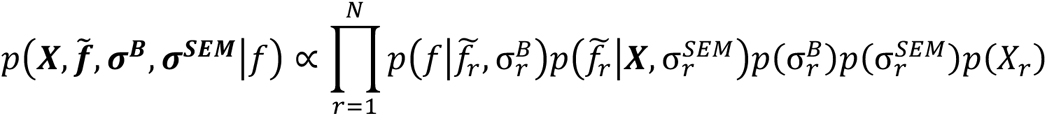

Where 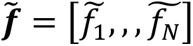 are the estimates of the contact frequencies made by each of the replicas r=1,…,N, determined using a forward model based on the chromatin conformations 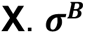 and 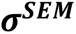 are respectively the systematic errors (from both forward model and experiments) and the standard errors of the mean due to the use of a finite number of replicas, respectively. The terms on the righthand side represent, for each replica, the probability of experimental contact frequency f given the forward model estimate 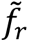, the probability of the estimate given the chromatin conformations **X**, the two prior probability distributions of the errors, and the prior probability of the chromatin conformation X_r_. For the forward model, we assume that the contact frequency (expressed as the number of sequencing reads) is estimated to be on average 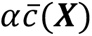, with α a proportionality constant to be determined, and the contact probability 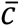 computed from an average over the replicas of the kernel function:

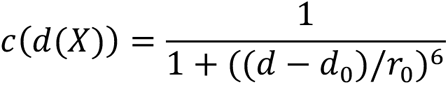

This function is close to 1 and 0 for short and large distances d between genomic loci, respectively, with d_0_ representing the distance at which the kernel is equal to 1, and r_0_ controlling its slope. Although these parameters depend on the experimental procedure and are in principle unknown, it is reasonable to assume that they should be close to the spatial resolution of the experiment, set by the typical distances over which cross-linking may occur. This kernel has been traditionally used to setup collective variables to explore protein folding with enhanced sampling all-atom MD simulations (72). Note that within this Bayesian framework, the accuracy of our forward model is accounted through the systematic errors 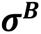.

Integrating the posterior over the estimated contact frequencies 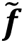 assuming Gaussian statistics and accounting for the entire LxL experimental contact map, we can sample the chromatin conformational ensemble by MD simulations using the posterior’s negative logarithm as an energy score (37):

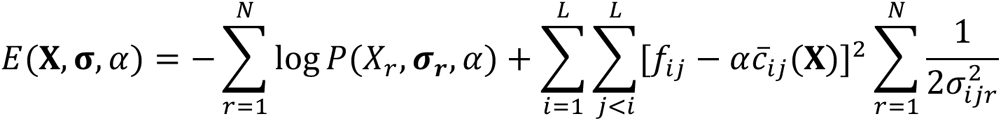

where the first term constitutes our prior, including the interaction potential of the original chromatin model, 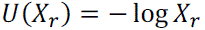, and the negative logarithm of the hyperparameter priors; f_ij_ and 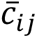 are respectively the experimental contact frequencies and the simulation contact probabilities between genomic locations i and j (the latter obtained by averaging the kernel function c(d_ij_) over all the replicas); 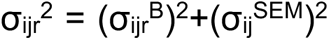 represents the total error on the i-j contact from replica r, resulting from the combination of systematic error (B) and standard error of the mean (SEM). Overall, the second term represents a harmonic bias forcing the system to adopt an ensemble of conformations whose observables are consistent with the experimental data. The standard errors of the means are estimated directly from the conformational ensemble, while systematic errors are sampled according to the posterior distribution using a noninformative prior via a Monte-Carlo scheme. The scaling factor α is also sampled by Monte-Carlo each step using a uniform prior. We implemented the Hi-C metainference protocol in PLUMED 2 (73) based on the existing metainference routines.

### Chromatin Models

In the 1CPN model (30), the nucleosome core particle is represented as a single ellipsoidal rigid body with orientation-dependent interactions with other nucleosomes, while linker DNA as a twistable wormlike chain (74) where each bead represents three base-pairs. Importantly, the 1CPN model can effectively account for spontaneous DNA sequence dependent nucleosome sliding (6), effectively allowing changes in linker DNA length. All 1CPN simulations have been performed using the software LAMMPS (75) at ambient temperature T=300K and 150 mM ionic strength, integrating the equations of motion by Langevin dynamics using a timestep of 60 fs and the default 1CPN settings (30). We set the DNA sequence parameter controlling nucleosome sliding to that of telomeric polyTTAGGG sequences, representing an average genomic sequence with intermediate nucleosome affinity.

In the 1kb model, chromatin is represented as a beads-on-a-string polymer model with interactions parametrized via a bottom-up procedure from 1CPN simulations of 50-nucleosome fibers with a nucleosome repeat length of 200bp, close to the average value observed in experiments of mESCs (67), our target system. The 1kb beads (each representing of 5 nucleosomes plus linker DNA) have a size of a=22nm, estimated from the average distance between the centers of mass of 1kb regions in the 1CPN simulations (Fig. 1C, left panel). The beads are linked along the polymer chain by harmonic bonds with potential E_b_(r)=12k_B_T(r/a-1)^2^ and by angle bonds E_a_(r)=k_B_T(cosθ+1)^2^ (with θ the angle formed by the central bead connecting its neighbors). The dihedral angle distribution obtained from 1CPN is uniform, so no extra dihedral potential was required. For non-local interactions, the 1kb beads interact with other non-neighboring beads via the Lennard-Jones potential E_nb_(r)=1.2k_B_T[(a/d)^12^-(a/d)^6^] within a distance cutoff of d_cut_=3a, where the strength has been set so that the polymer size scales as in the theta condition (size∼L^½^, Fig. S1D), at which the conformational ensemble includes both compact and open conformations. The 1CPN statistics used to parametrize our model has been produced from 32 trajectories of 4×10^9^ timesteps, each equilibrated for 5×10^9^ steps starting from extended conformations (see Fig. S1A-C for the equilibration of radius of gyration, neighboring distances and angles). During model parametrization, the unbiased 1kb simulations have been performed using LAMMPS (75) at ambient temperature T=300K, integrating the equations of motion by Langevin dynamics using a timestep 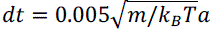, where the mass of the 1kb beads m is set to the default mass unit of 1 g/mole. Within the simulation, we set the bead sizes to the default LAMMPS unit of 1Å, but this is later rescaled to a=22nm when measuring actual physical distances. For the model parametrization, the bond and angle potential strengths were determined directly from the 1CPN distributions by Boltzmann inversion, while the non-bonded interaction strength for theta condition was determined from 10^7^-steps simulations with various strengths for polymer length of 50, 100, 200, 400, and 800 kb (Fig. S1D).

In the copolymer model used for Hi-C metainference validation, the three sticky CREs are those with bead ids 31-40 (E1), 71-80 (E2) and 151-160 (E3). In this copolymer, all non-neighboring beads interact via a Lennard-Jones potential E_nb_(r)=ε[(a/d)^12^-(a/d)^6^] for d<d_cut_=3a, with strength ε=3.2k_B_T for CRE-CRE pairs and ε=1.2k_B_T for any other pair (as in our prior). The target copolymer ensemble was generated from a long simulation (10^8^-steps) in the absence of metainference bias, from which we constructed a synthetic contact map using the kernel c_ij_=1/[1+((d_ij_-d_0_)/r_0_)^6^] with r_0_=a=22nm and d_0_=0.5a, and a scale factor of α=50 (meaning that a contact probability of 1 will correspond to 50 Hi-C reads).

### Hi-C Metainference Simulations

All the Hi-C metainference simulations were performed using the MD software LAMMPS (75) patched with PLUMED 2 (73) implementing Hi-C metainference. All simulations using the 1CPN and 1kb prior models were run using the same MD parameters used for the unbiased simulations described above. In our applications to the 3D genomic organization of mESCs, we augmented our 1CPN and 1kb chromatin models by metainference MD using the normalized Micro-C contact frequencies (39) of the target regions, at a resolution of either 200bp or 1kb (for the 1CPN and 1kb models, respectively). Raw data were processed using the software Juicebox (76). For the forward model used to estimate the contact frequency from the chromatin ensemble, we use a size parameter of r_0_=13nm and r_0_=a=22nm for the 1CPN and the 1kb simulations respectively, with these values corresponding to roughly the size of a nucleosome and that of a 1kb bead, a choice consistent with the resolution of the respective models and data. In both cases we set the kernel shift parameter as d_0_=r_0_/2, making the kernel close to the attractive part of a Lennard-Jones potential. When using the 1kb model, the metainference biasing force originating from a given contact frequency between two genomic locations is applied to the two corresponding 1kb beads, while for the 1CPN simulations the bias is applied to the corresponding nucleosome core particles. Metainference allows a variety of error models and priors, and upon evaluation of convergence speed and accuracy, we chose to use a single Gaussian error model acting on all the entries of the target contact map. For all mESC applications, we ran simulations using 128 replicas. When integrating the Hi-C map into MD, we also include in the metainference bias distant loci with zero Hi-C reads (so pairs of loci where contacts are not observed in experiments), unless they correspond to a genomic location where all the entries are zero (which likely reflects issues with sequencing at this location).

We applied metainference to reconstruct the 3D organization of the Pou5f1 (chr 17: 35,420-35,720kb), Sox2 (chr 3: 34,620-34,920kb), and Nanog (chr 6: 122,600-122,800kb) genomic loci in mESCs (mm10). These genomic loci have been often analyzed in single cell experiments focusing on gene regulation (46, 77), aiding the validation of our simulations and the discussion on the possible interplay between gene structure and function. All these loci were first studied at 1kb resolution by running 10^6^-steps Hi-C metainference simulations (the first 10^5^ steps of each simulation were discarded as equilibration). We then employed 1CPN (30) metainference simulations to explore in more detail the organization of a 10kb Nanog region (chr 6: 122,701-122,711kb) encompassing the promoter (chr 6: 122,707kb) and its closest enhancer located 5kb upstream. For these simulations, we chose the initial chromatin conformations among those sampled during the unbiased 1CPN trajectories, selecting for each of the 128 replicas the 10kb conformation with the smallest root mean square distance to the corresponding one observed at the last step of the 1kb metainference simulations (this ensures that the initial 1CPN ensemble is already in reasonable agreement with the target Micro-C map). The 1CPN Nanog simulation was 6×10^6^ MD steps long and we discarded the first 5×10^6^ steps as equilibration.

For comparison with real timescales, we rescaled the simulation time of the 1kb model simulations so that the diffusion of chromatin within the Sox2 genomic locus matches the experimental estimate of 0.25 μm^2^/s (47) (Fig. S3B).

## Data Availability

Data to reproduce the reported analysis and code implementing Hi-C metainference will be made available upon publication.

## References

1. E. Chacin, et al., Establishment and function of chromatin organization at replication origins. Nature 2023, 1–7 (2023).

2. B. Bonev, G. Cavalli, Organization and function of the 3D genome. Nat Rev Genet 17, 661– 678 (2016).

3. E. H. Finn, T. Misteli, Molecular basis and biological function of variability in spatial genome organization. Science (1979) 365, eaaw9498 (2019).

4. K. Luger, A. W. Mäder, R. K. Richmond, D. F. Sargent, T. J. Richmond, Crystal structure of the nucleosome core particle at 2.8 Å resolution. Nature 389, 251–260 (1997).

5. K. Zhou, G. Gaullier, K. Luger, Nucleosome structure and dynamics are coming of age. Nat Struct Mol Biol 26, 3–13 (2019).

6. G. B. Brandani, T. Niina, C. Tan, S. Takada, DNA sliding in nucleosomes via twist defect propagation revealed by molecular simulations. Nucleic Acids Res. 46, 2788–2801 (2018).

7. C. Tan, S. Takada, Nucleosome allostery in pioneer transcription factor binding. Proc. Natl. Acad. Sci. U.S.A. 117, 20586–20596 (2020).

8. C. M. Maccarthy, et al., OCT4 interprets and enhances nucleosome flexibility. Nucleic Acids Res 50, 10311–10327 (2022).

9. A. Soman, et al., Columnar structure of human telomeric chromatin. Nature 609, 1048–1055 (2022).

10. T. Schalch, S. Duda, D. F. Sargent, T. J. Richmond, X-ray structure of a tetranucleosome and its implications for the chromatin fibre. Nature 436, 138–141 (2005).

11. B. Ekundayo, T. J. Richmond, T. Schalch, Capturing Structural Heterogeneity in Chromatin Fibers. J Mol Biol 429, 3031–3042 (2017).

12. Z. Hou, F. Nightingale, Y. Zhu, C. MacGregor-Chatwin, P. Zhang, Structure of native chromatin fibres revealed by Cryo-ET in situ. Nat Commun 14, 6324 (2023).

13. M. Ohno, et al., Sub-nucleosomal Genome Structure Reveals Distinct Nucleosome Folding Motifs. Cell 176, 520–534 (2019).

14. J. R. Dixon, et al., Topological domains in mammalian genomes identified by analysis of chromatin interactions. Nature 485, 376–380 (2012).

15. G. R. Cavalheiro, T. Pollex, E. E. Furlong, To loop or not to loop: what is the role of TADs in enhancer function and gene regulation? Curr Opin Genet Dev 67, 119–129 (2021).

16. M. Yokoshi, K. Segawa, T. Fukaya, Visualizing the Role of Boundary Elements in Enhancer-Promoter Communication. Mol Cell 78, 224–235.e5 (2020).

17. J. Zuin, et al., Nonlinear control of transcription through enhancer-promoter interactions. Nature 604 (2022).

18. D. B. Brückner, H. Chen, L. Barinov, B. Zoller, T. Gregor, Stochastic motion and transcriptional dynamics of pairs of distal DNA loci on a compacted chromosome. Science 380, 1357–1362 (2023).

19. M. V. Imakaev, G. Fudenberg, L. A. Mirny, Modeling chromosomes: Beyond pretty pictures. FEBS Lett 589, 3031–3036 (2015).

20. L. Giorgetti, et al., Predictive polymer modeling reveals coupled fluctuations in chromosome conformation and transcription. Cell 157, 950–63 (2014).

21. M. Di Pierro, B. Zhang, E. L. Aiden, P. G. Wolynes, J. N. Onuchic, Transferable model for chromosome architecture. Proc Natl Acad Sci U S A 113, 12168–12173 (2016).

22. S. Shinkai, et al., PHi-C: deciphering Hi-C data into polymer dynamics. NAR Genom Bioinform 2 (2020).

23. G. Shi, D. Thirumalai, From Hi-C Contact Map to Three-Dimensional Organization of Interphase Human Chromosomes. Phys Rev X 11, 011051 (2021).

24. E. Lieberman-Aiden, et al., Comprehensive mapping of long-range interactions reveals folding principles of the human genome. Science 326, 289–93 (2009).

25. G. Fudenberg, et al., Formation of Chromosomal Domains by Loop Extrusion. Cell Rep 15, 2038–2049 (2016).

26. C. A. Brackley, et al., Predicting the three-dimensional folding of cis-regulatory regions in mammalian genomes using bioinformatic data and polymer models. Genome Biol 17, 59 (2016).

27. D. Jost, P. Carrivain, G. Cavalli, C. Vaillant, Modeling epigenome folding: formation and dynamics of topologically associated chromatin domains. Nucleic Acids Res 42, 9553–61 (2014).

28. Q. MacPherson, B. Beltran, A. J. Spakowitz, Bottom-up modeling of chromatin segregation due to epigenetic modifications. Proc Natl Acad Sci U S A 115, 12739–12744 (2018).

29. G. Arya, Q. Zhang, T. Schlick, Flexible histone tails in a new mesoscopic oligonucleosome model. Biophys J 91, 133–150 (2006).

30. J. Lequieu, A. Córdoba, J. Moller, J. J. De Pablo, 1CPN: A coarse-grained multi-scale model of chromatin. Journal of Chemical Physics 150, 215102 (2019).

31. S. Portillo-Ledesma, Z. Li, T. Schlick, Genome modeling: From chromatin fibers to genes. Curr Opin Struct Biol 78 (2023).

32. S. E. Farr, E. J. Woods, J. A. Joseph, A. Garaizar, R. Collepardo-Guevara, Nucleosome plasticity is a critical element of chromatin liquid–liquid phase separation and multivalent nucleosome interactions. Nat. Commun. 12, 1–17 (2021).

33. G. D. Bascom, C. G. Myers, T. Schlick, Mesoscale modeling reveals formation of an epigenetically driven HOXC gene hub. Proceedings of the National Academy of Sciences 116, 4955–4962 (2019).

34. K. Shrinivas, et al., Enhancer Features that Drive Formation of Transcriptional Condensates. Mol Cell 75, 549–561.e7 (2019).

35. T.-H. S. Hsieh, et al., Mapping Nucleosome Resolution Chromosome Folding in Yeast by Micro-C. Cell 162, 108–119 (2015).

36. S. Carstens, M. Nilges, M. Habeck, Bayesian inference of chromatin structure ensembles from population-averaged contact data. Proc Natl Acad Sci U S A 117, 201910364 (2020).

37. M. Bonomi, C. Camilloni, A. Cavalli, M. Vendruscolo, Metainference: A Bayesian inference method for heterogeneous systems. Sci Adv 2, e1501177 (2016).

38. J. Moller, J. Lequieu, J. J. De Pablo, The Free Energy Landscape of Internucleosome Interactions and Its Relation to Chromatin Fiber Structure. ACS Cent. Sci. 5, 341–348 (2019).

39. T. H. S. Hsieh, et al., Resolving the 3D Landscape of Transcription-Linked Mammalian Chromatin Folding. Mol Cell 78, 539–553.e8 (2020).

40. A. Grosberg, Y. Rabin, S. Havlin, A. Neer, Crumpled globule model of the three-dimensional structure of dna. Europhys Lett 23, 373–378 (1993).

41. B. Bintu, et al., Super-resolution chromatin tracing reveals domains and cooperative interactions in single cells. Science 362, eaau1783 (2018).

42. B. A. Gibson, et al., Organization of Chromatin by Intrinsic and Regulated Phase Separation. Cell 179, 470–484.e21 (2019).

43. C. M. Bishop, Pattern Recognition and Machine Learning (Springer, 2006).

44. Y. Naritomi, S. Fuchigami, Slow dynamics in protein fluctuations revealed by time-structure based independent component analysis: The case of domain motions. J Chem Phys 134, 065101 (2011).

45. M. K. Scherer, et al., PyEMMA 2: A Software Package for Estimation, Validation, and Analysis of Markov Models. J Chem Theory Comput 11, 5525–5542 (2015).

46. J. Li, et al., Single-gene imaging links genome topology, promoter–enhancer communication and transcription control. Nat Struct Mol Biol 27, 1032–1040 (2020).

47. C. H. Bohrer, D. R. Larson, Synthetic analysis of chromatin tracing and live-cell imaging indicates perva-sive spatial coupling between genes. Elife (2023) 10.1101/2022.07.07.499202 (March 7, 2023).

48. J. Rodriguez, D. R. Larson, Annual Review of Biochemistry Transcription in Living Cells: Molecular Mechanisms of Bursting. Annu Rev Biochem 89, 189–212 (2020).

49. T. Nozaki, et al., Condensed but liquid-like domain organization of active chromatin regions in living human cells. Sci Adv 9, eadf1488 (2023).

50. S. Blinka, S. Rao, Nanog Expression in Embryonic Stem Cells – An Ideal Model System to Dissect Enhancer Function. BioEssays 39, 1700086 (2017).

51. I. Zhu, W. Song, I. Ovcharenko, D. Landsman, A model of active transcription hubs that unifies the roles of active promoters and enhancers. Nucleic Acids Res 49, 4493–4505 (2021).

52. M. H. Kagey, et al., Mediator and cohesin connect gene expression and chromatin architecture. Nature 467, 430–435 (2010).

53. C. Chronis, et al., Cooperative Binding of Transcription Factors Orchestrates Reprogramming. Cell 168, 442–459.e20 (2017).

54. J. A. Stamatoyannopoulos, et al., An encyclopedia of mouse DNA elements (Mouse ENCODE). Genome Biol 13, 1–5 (2012).

55. T. Brouwer, et al., A critical role for linker DNA in higher-order folding of chromatin fibers. Nucleic Acids Res 49, 2537–2551 (2021).

56. H. Ishii, J. T. Kadonaga, B. Ren, MPE-seq, a new method for the genome-wide analysis of chromatin structure. Proc Natl Acad Sci U S A 112, E3457–E3465 (2015).

57. W. F. Richter, S. Nayak, J. Iwasa, D. J. Taatjes, The Mediator complex as a master regulator of transcription by RNA polymerase II. Nat Rev Mol Cell Biol 23, 732–749 (2022).

58. D. Krepel, R. R. Cheng, M. Di Pierro, J. N. Onuchic, Deciphering the structure of the condensin protein complex. Proc Natl Acad Sci U S A 115, 11911–11916 (2018).

59. G. Forte, et al., Transcription modulates chromatin dynamics and locus configuration sampling. Nat Struct Mol Biol 30, 1275–1285 (2023).

60. A. Panigrahi, B. W. O’Malley, Mechanisms of enhancer action: the known and the unknown. Genome Biol 22 (2021).

61. N. S. Benabdallah, et al., Decreased Enhancer-Promoter Proximity Accompanying Enhancer Activation. Mol Cell 76, 473–484.e7 (2019).

62. J. M. Alexander, et al., Live-cell imaging reveals enhancer-dependent Sox2 transcription in the absence of enhancer proximity. Elife 8 (2019).

63. J. P. Karr, J. J. Ferrie, R. Tjian, X. Darzacq, The transcription factor activity gradient (TAG) model: contemplating a contact-independent mechanism for enhancer–promoter communication. Genes Dev 36, 7–16 (2022).

64. J. Xiao, A. Hafner, A. N. Boettiger, How subtle changes in 3d structure can create large changes in transcription. Elife 10 (2021).

65. R. Collepardo-Guevara, et al., Chromatin Unfolding by Epigenetic Modifications Explained by Dramatic Impairment of Internucleosome Interactions: A Multiscale Computational Study. J Am Chem Soc 137, 10205–10215 (2015).

66. R. Zhang, J. Erler, J. Langowski, Histone Acetylation Regulates Chromatin Accessibility: Role of H4K16 in Inter-nucleosome Interaction. Biophysical J. 112, 450–459 (2017).

67. L. N. Voong, et al., Insights into Nucleosome Organization in Mouse Embryonic Stem Cells through Chemical Mapping Article Insights into Nucleosome Organization in Mouse Embryonic Stem Cells through Chemical Mapping. Cell 167, 1555–1570.e15 (2016).

68. M. Gabriele, et al., Dynamics of CTCF– and cohesin-mediated chromatin looping revealed by live-cell imaging. Science (1979) 376, 476–501 (2022).

69. G. Mohana, et al., Chromosome-level organization of the regulatory genome in the Drosophila nervous system. Cell 186, 3826–3844.e26 (2023).

70. K. Maeshima, S. Iida, M. A. Shimazoe, S. Tamura, S. Ide, Is euchromatin really open in the cell? Trends Cell Biol (2023) 10.1016/j.tcb.2023.05.007.

71. B. A. Gibson, et al., In diverse conditions, intrinsic chromatin condensates have liquid-like material properties. Proc Natl Acad Sci U S A 120 (2023).

72. S. Piana, A. Laio, A bias-exchange approach to protein folding. J Phys Chem B 111, 4553–9 (2007).

73. G. A. Tribello, M. Bonomi, D. Branduardi, C. Camilloni, G. Bussi, PLUMED 2: New feathers for an old bird. Comput Phys Commun 185, 604–613 (2014).

74. C. A. Brackley, A. N. Morozov, D. Marenduzzo, Models for twistable elastic polymers in Brownian dynamics, and their implementation for LAMMPS. J Chem Phys 140, 135103 (2014).

75. S. Plimpton, Fast parallel algorithms for short-range molecular dynamics. J Comput Phys 117, 1–19 (1995).

76. J. T. Robinson, et al., Juicebox.js Provides a Cloud-Based Visualization System for Hi-C Data. Cell Syst 6, 256–258.e1 (2018).

77. H. Ohishi, et al., STREAMING-tag system reveals spatiotemporal relationships between transcriptional regulatory factors and transcriptional activity. Nat Commun 13, 7672 (2022).

