## Supporting Information for "Multiscale Bayesian Simulations Reveal Functional Chromatin Condensation of Gene Loci"

### Analysis of the copolymer trajectories by Markov state modeling

To analyze the conformational ensembles and the dynamics of the polymers for our Hi-C metainference validation, we first performed TICA dimensionality reduction (1) on the copolymer contact maps defined from the positions of every $5^{th}$ 1kb beads using the kernel $c_{ij}(d_{ij})=1/(1+d_{ij}/r_0)$ with $r_0$=22 nm. The coordinates of the polymers in our Hi-C metainference simulations have been transformed using the same linear TICA transformation determined from the copolymer simulations. Fig. 2D and SI Fig. 2D show the free energy landscapes along the two slowest TICA coordinates for the Hi-C metainference (128 replicas) and copolymer MD simulations, respectively. To quantitatively compare the probability distributions of the metainference and target copolymer ensembles, we divided the ensembles into 50 clusters by k-means clustering (2) using the 6 slowest TICA coordinates of the copolymer system. The Hi-C metainference conformations have been equivalently divided into 50 clusters based on the same cluster centers determined from the copolymer clustering. Fig. 2E indicates the Kullback-Leibler between the metainference and copolymer equilibrium probability distributions of these 50 clusters. To evaluate the dynamics, we estimated, for the metainference and copolymer systems, the transitions probabilities between the clusters by Markov state modeling using a lag-time of $10^4$ MD steps (3) (Fig. 2G from main text).

### Supplementary figures

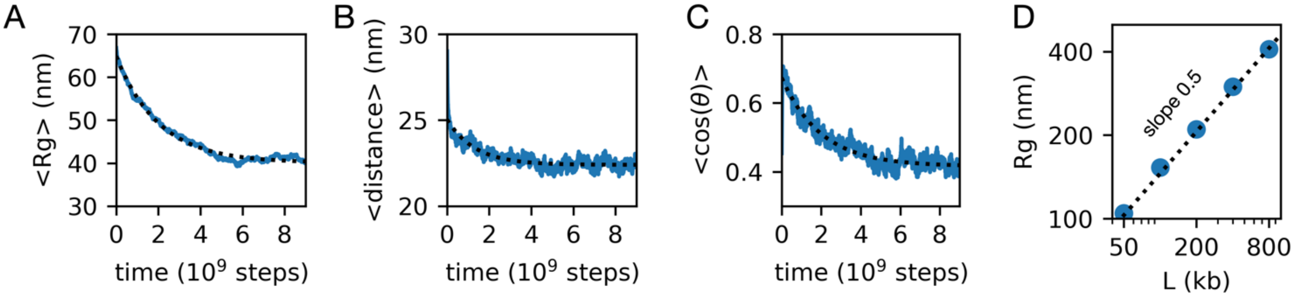


Figure S1. Unbiased chromatin fiber MD simulations. A-C. Running averages over 32 independent 50-nucleosome 1CPN trajectories showing the equilibration of radius of gyration (A), and chromatin fiber neighbor distances (B) and cosine of the angles (C) over time. To compute the distances and angles, the chromatin fiber is coarse-grained to 1kb beads, each located at the center of mass of 5 neighboring nucleosomes. The neighbor distances and angles are calculated based on the positions of these 1kb beads along the fiber (as indicated in Fig. 1A from the main text). For the parametrization of the 1kb model, the first $5 \times 10^9$ MD steps are discarded as equilibration. The dotted lines are exponential fits to the data. D. Scaling of the average radius of gyration (Rg) with chromatin fiber length in our 1kb resolution model, plotted on a logarithmic scale, based on $10^7$-steps MD simulations. The slope of 0.5 indicates that the radius of gyration grows as $Rg \sim L^{0.5}$, as an ideal chain.

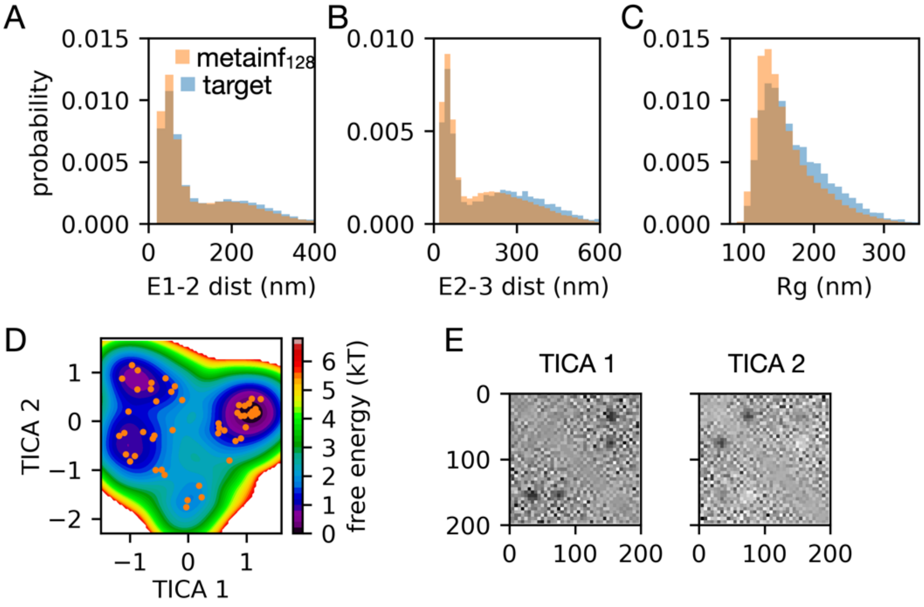


Figure S2. Hi-C metainference validation. A-C. Comparison between metainference (128 replicas) and target copolymer ensemble probability distributions of the distances between enhancer-like elements 1 and 2 (A) and 2 and 3 (B), and the radius of gyration (C). D. Free energy landscape of the target copolymer ensemble obtained by projecting the conformations along the two slowest TICA coordinates. E. Components of the two slowest TICA coordinates, capturing different interactions between the three enhancer-like elements of the copolymer.

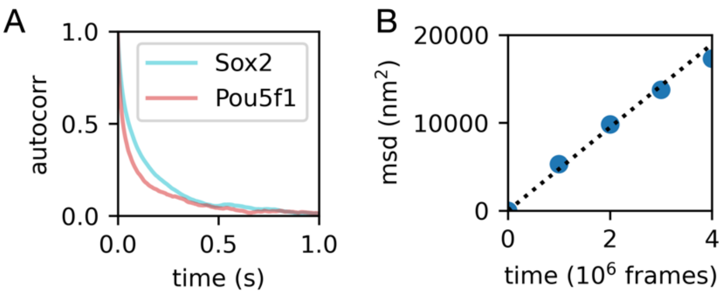


Figure S3. Hi-C metainference of the Sox2 and Pou5f1 loci of mESCs using the 1kb model. A. Time autocorrelations of the distances ($<\Delta d(t)\Delta d(t+\tau)>_t/<\Delta d^2>$) between the Sox2 and Pou5f1 promoters and their respective enhancers +100kb and +40kb away. B. Mean square displacement of the Sox2 enhancer-promoter 3d distance $<[\Delta \mathbf{r}(t+\tau)-\Delta \mathbf{r}(t)]^2>_t$ as a function of simulation timestep. The scaling factor to convert simulation time to real time (reported throughout the main text) is obtained by matching the typical chromatin experimental diffusion coefficient of 0.25 μm$^2$/s (4) based on a linear fit within a small time window (dotted line). TODO add comparison of Sox2 and Pou5f1 contact probability between metainference and experiments.

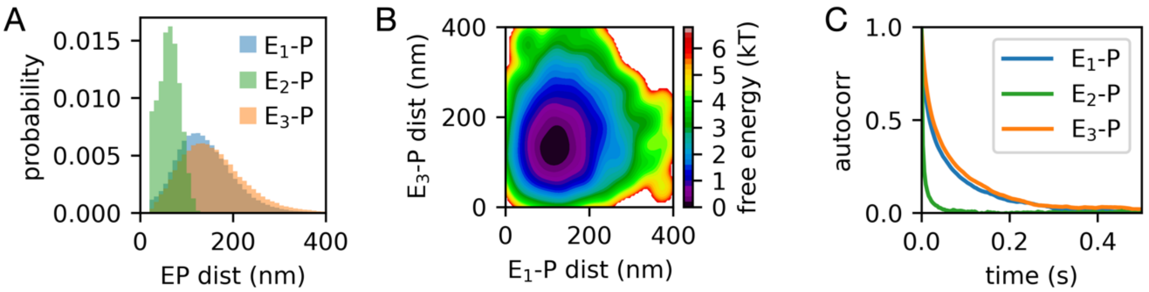


Figure S4. Hi-C metainference of the Nanog locus of mESCs using the 1kb model. A. Probability distributions of the distances between the Nanog promoter and its three enhancers located at -40kb ($E_1$), -5kb ($E_2$), and +60kb ($E_3$) positions. B. Free energy landscape along the distances between the Nanog promoter and the -40kb ($E_1$) and +60kb ($E_3$) enhancers. C. Time autocorrelations of the distances ($<\Delta d(t)\Delta d(t+\tau)>_t/<\Delta d^2>$) between the Nanog promoter and its three enhancers at -40kb ($E_1$), -5kb ($E_2$), and +60kb ($E_3$) positions.

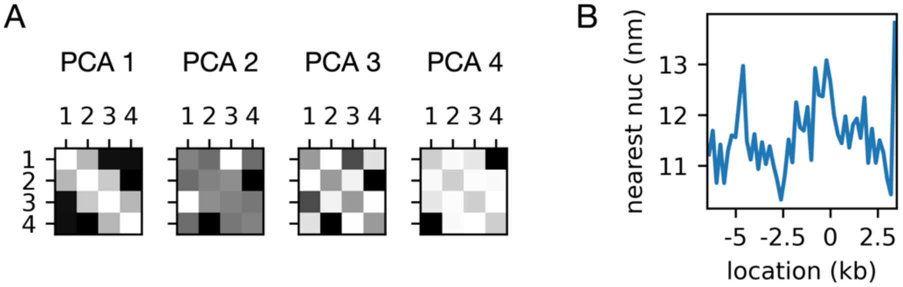


Figure S5. A. Eigenvectors of the first four tetranucleosome principal components, projected on the tetranucleosome contact maps (the coordinates used for PCA). Darker contacts correspond to higher eigenvector components. B. For each nucleosome along the 10kb Nanog locus (x-axis), the average distance to the nearest nucleosome as observed in the 1CPN metainference ensemble.

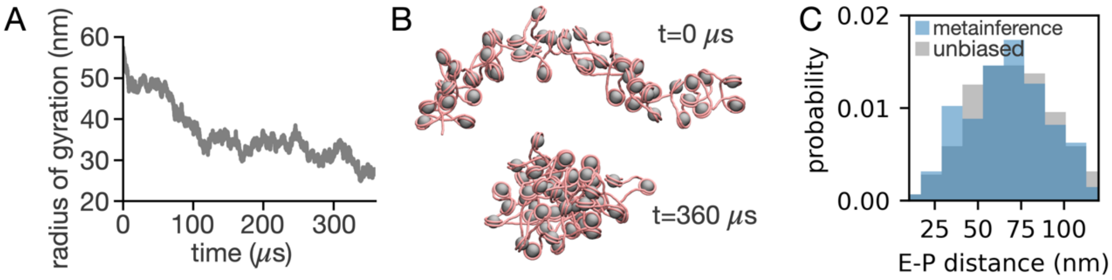


Figure S6. A. Representative 360-$\mu$s unbiased MD trajectory using the 1CPN prior model (in the absence of metainference). B. Snapshots of the initial and final chromatin configurations of the trajectory in panel A. C. Comparison of the probability distributions of the enhancer-promoter distances observed in the Hi-C metainference (blue) and unbiased (grey) MD simulations. The unbiased ensemble was generated from the same 36 4x$10^9$-steps trajectories (excluding equilibration) used to parametrize the 1kb prior model of chromatin (these are also described in SI Fig. 1). The prior 1CPN system is made of 50 nucleosomes with a uniform 200 bp nucleosome repeat region; here the enhancer-promoter distances represent the distances between the nucleosomes closest to the corresponding enhancer and promoter regions in the Hi-C metainference simulations.

### References


1. Y. Naritomi, S. Fuchigami, Slow dynamics in protein fluctuations revealed by time-structure based independent component analysis: The case of domain motions. *J Chem Phys* **134**, 065101 (2011).
2. C. M. Bishop, *Pattern Recognition and Machine Learning* (Springer, 2006).
3. M. K. Scherer, *et al.*, PyEMMA 2: A Software Package for Estimation, Validation, and Analysis of Markov Models. *J Chem Theory Comput* **11**, 5525–5542 (2015).
4. C. H. Bohrer, D. R. Larson, Synthetic analysis of chromatin tracing and live-cell imaging indicates perva-sive spatial coupling between genes. *Elife* (2023) https:/doi.org/10.1101/2022.07.07.499202 (March 7, 2023).
